# Humans and marmosets share similar face recognition signatures in shape-based visual face discrimination behavior

**DOI:** 10.1101/2024.11.19.624374

**Authors:** You-Nah Jeon, Hector Y. Cho, Ashley C. Green, Elias B. Issa

## Abstract

Our ability to identify faces is thought to depend on high-level visual processing in the brain. Nonetheless, studies of face recognition have generally relied on 2D face photographs where low-level strategies relying on texture and appearance cues can be employed to adequately support high face identification performance. Here, we designed a fine face discrimination task under 3D pose and lighting variation that was purely based on shape, a task which challenged state-of-the-art artificial vision systems compared to object recognition tasks. In contrast, humans performed this shape based face task at comparable levels to their object recognition performance. We then tested one of the smallest simian primates on this human-level, machine-difficult visual task, the common marmoset – a small, New World monkey. Marmosets successfully discriminated between face identities across 3D viewing conditions based purely on face shape. Their face recognition performance was on par with their object recognition performance and exhibited face-specific behavioral signatures similar to humans, including lower performance for inverted faces, faces lit from below, and contrast reversed faces. These results demonstrate that a high-level visual behavior, invariant face recognition based purely on geometry and not additional texture and appearance cues, is shared across simian primates from among the smallest to the most advanced, consistent with the presence of common underlying high-level visual brain areas across simian primates.

## INTRODUCTION

Face perception is thought to depend on several hierarchically organized cortical areas in the ventral visual pathway,^1,2^ but defining a task that specifically engages such high-level face processing remains an open challenge. Previous work on invariant object recognition (also known as core object recognition) emphasized the importance of testing the ability to recognize objects across many identity-preserving transformations, including their position, scale (i.e. distance from the observer), background, and clutter^3–5^. Stimulus sets created with such variation have been used to compare different biological visual systems – human, macaque, marmoset – to each other and to artificial visual systems^6–10^. Compared to these object recognition studies, face perception research, whether behavioral or neural, whether human, monkey or artificial, has typically utilized photographs of faces or synthetic faces of either frontal or very limited views, of similar illumination, of similar size, and often without rich, varied backgrounds^11–21^. These stimuli, such as face photographs in the wild are seemingly natural but lack adequate independently controlled variations. The multiple cues that are present in these stimuli permit use of conflating low-level features to solve the identification task and relatedly, obfuscate what cortical mechanisms are necessary for robustly performing the behavior^22^. For example, various textural cues such as individual-specific hair or eye color in photographs can be easily picked up by simple visual systems, but the recognition performance of low-level visual features typically does not generalize to identify the same individuals in novel settings.

Here, we created a novel face discrimination task – with textureless face images generated purely based on the geometry of the face – to rigorously examine shape-based face recognition behavior in humans and monkeys. Moreover, we incorporated greater identity-preserving transformations, namely face pose, lighting direction, and background variation, than previous face recognition studies. Remarkably, the resulting task was found to be much more challenging for artificial vision systems than the original core object recognition task, suggesting the specific challenge posed by the finer shape discrimination required for faces versus objects. We found that both humans and marmosets robustly performed the challenging face discrimination task while also displaying comparable general behavioral signatures of face recognition. Our results suggest that particular aspects of high-level face perception are shared across the simian primate clade, complementing the growing neuroscientific findings on the functional and structural homology of the ventral visual stream across marmosets, macaques, and humans^18,19,23,24^.

## RESULTS

### Humans and marmosets recognized faces defined by shape across changes in rotation, scale, lighting direction, and background

We evaluated human and marmoset ability to discriminate geometrically defined faces invariant to identity-preserving changes in rotation, scale, lighting direction and background. Our stimulus set included an equal number of upright and inverted faces rendered across three different lighting contexts: lighting from above (i.e. typical lighting), lighting from below, and contrast reversal (**Figure 1**). Subjects, whether human or marmoset, participated in a two-alternative forced choice task (2AFC) where they chose which of the two identities was present in the flashed image (**Figure 2A**; see **Video S1** of marmoset performing the 2AFC task). To benchmark the difficulty of this face discrimination task and ensure it was not solvable by low-level features, we used a state-of-the-art deep convolutional neural network (DNN) model, ResNet-50^28^, pretrained on the ImageNet database. DNNs trained to categorize images broadly match human image categorization performance^26^ and are considered the leading models of human and macaque ventral visual cortex^6,7,27,28^. While an ImageNet-pretrained ResNet-50 achieved 92.5% on the 2AFC object discrimination task previously studied in the lab^10^, comparable to human performance levels, the same model only managed 59% performance on the 2AFC face task (chance=50%), exhibiting a steep fall-off in performance from the object recognition task (**Figure 2B**; compare red to gray bars). This task-specific drop in model performance occurred even though the object task images were rendered in a similar fashion to the face task images – both tasks utilized grayscale renderings from synthetic 3D meshes superimposed on random 2D natural image backgrounds. The poor face task performance was not remedied by the number of face images used for training the linear SVM decoder nor by using a DNN encoder explicitly trained to categorize faces from a large corpus of face photographs (VGG-Face) (**Figure S1**). That face-trained network performed at 98.95% on face photographs but only 57% on our binary shape-based face discrimination task^29^.

**Figure 1.**
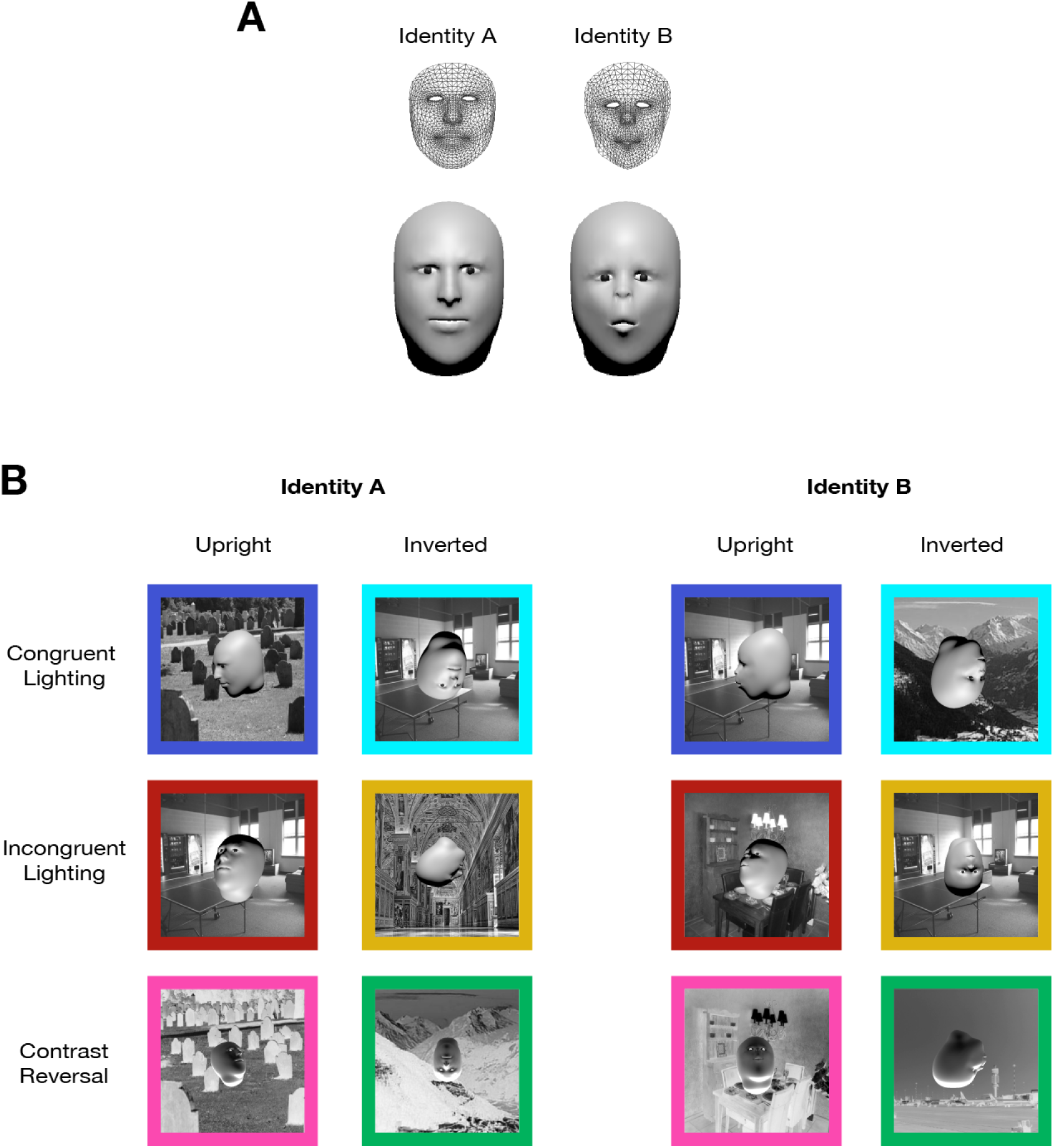
Face Stimuli. **A *Two face identities.*** Original meshes acquired using Apple Face Capture depicted as wireframes and the output renderings from the final textureless 3D face models with eyes and head added. **B *Face orientation and lighting conditions.*** We combined two in-plane rotations, upright and inverted, and three lighting contexts, lighting from above, lighting from below, and contrast reversal, to create six different conditions. Congruent lighting refers to the condition where the lighting source is overhead. An example image from each of the six different conditions is shown for both identities (25 images per condition used in the task).

**Figure 2.**
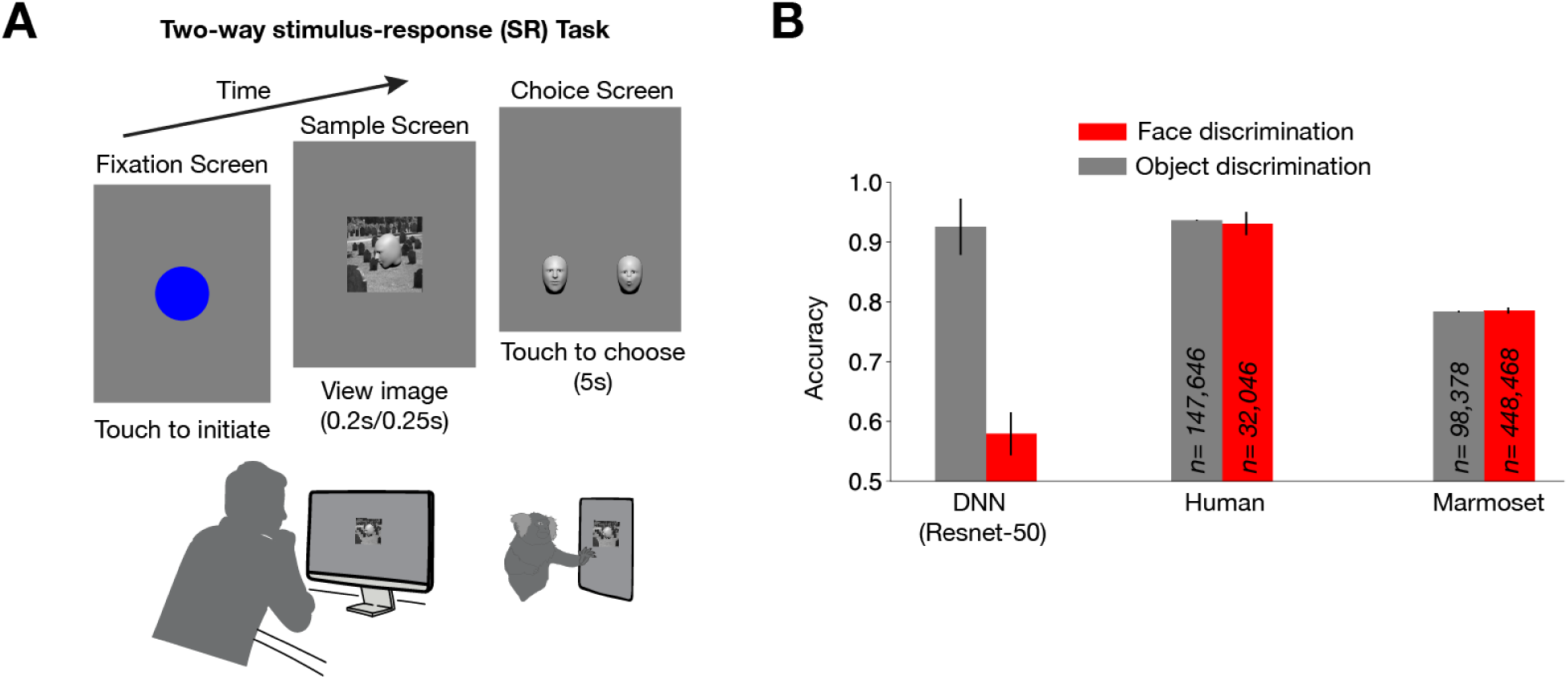
Face discrimination task and performance. A *Task design*. Both humans and marmosets performed a 2-way stimulus-response task where they chose which of the two identities a sample image corresponded to. The sample image was shown for 250ms for marmosets and 200ms for humans. **B *Task performance.*** Humans and marmosets far outperformed a DNN (ResNet-50) on the face discrimination task. For comparison, performance numbers for the object discrimination task (50ms for humans and 250ms for marmosets) were adapted from *Kell et al*. Error bars indicate standard deviations across bootstrap resamples over trials for human and marmoset. Error bars for ResNet-50 are standard deviations across bootstrap resamples of training images.

Far exceeding the performance of DNNs, humans performed our 3D face task at 91% (pooled across 4 human subjects; individual subjects performed 88%, 97%, 94%, and 85%; See **Methods** for more details on human subject pools), and marmosets performed at 78% (4 marmosets pooled; individual subjects performed 71%, 82%, 80%, and 76%; n=1863 trials/session, **Figure S2**). Notably, both humans and marmosets did not show any appreciable performance drop-off on the face task compared to their performance on the object discrimination task (**Figure 2B**; compare red to gray bars). For all human and marmoset performance numbers, we applied a performance correction based on the lapse rate calculated as the subject’s error rate for the best image. Both humans and marmosets had a lapse rate of ∼2%. This mild 2% adjustment allowed for quantitative comparisons of absolute visual ability (*sans* lapses in attention and/or downstream motor response errors during some trials). Thus, lapse-corrected primate performance allows direct comparison to DNNs which effectively have zero lapse rate as decoding of the face identity was done directly from sensory representations in artificial vision models, without additional sources of attentional or motor noise as in the biological system.

### Face inversion and contrast reversal deficits in humans during shape based, fine face discrimination

To test whether our face discrimination task tapped into canonical face processing mechanisms, we assessed two well-known observations from human face behavior: the inversion ^30,31^ and contrast reversal effects^32,33^. Prior work had demonstrated these effects in a different task setting than ours, so we sought to determine whether inversion and contrast reversal effects manifest in performing a shaped-based face discrimination task. Because the face inversion effect is meaningful when quantitatively larger than the size of the inversion effect for objects, we prepared an object discrimination task for comparison to the face task using the same distributions for each latent parameter – rotation, scale, lighting, and background **(Figure 3A)**. We chose camel and elephant meshes as two object identities which, like faces, have a stereotyped configuration, namely a head at the top, a body, and four legs below^34^. Again, to focus on fine geometry discrimination, we rendered the camel and elephant meshes without texture. When we compared performance for the inverted versus upright conditions in the two tasks (camel vs. elephant and face identity A vs. face identity B), we found a disproportionately stronger decrease in inverted versus upright performance for the face task compared to the object task (two tailed t-test: t = −14.30, p = 7.16 × *e*^−17^; **Figure 3B**). Our tasks also revealed that contrast reversal induced a performance deficit in the face task that was stronger than when contrast reversal was applied in the setting of shape-based object discrimination (two tailed t-test: t = −3.35, p = 0.0018; **Figure 3C**). These results confirmed that our shape-based fine face discrimination paradigm evoked face-specific visual processing deficits similar to prior work.

**Figure 3.**
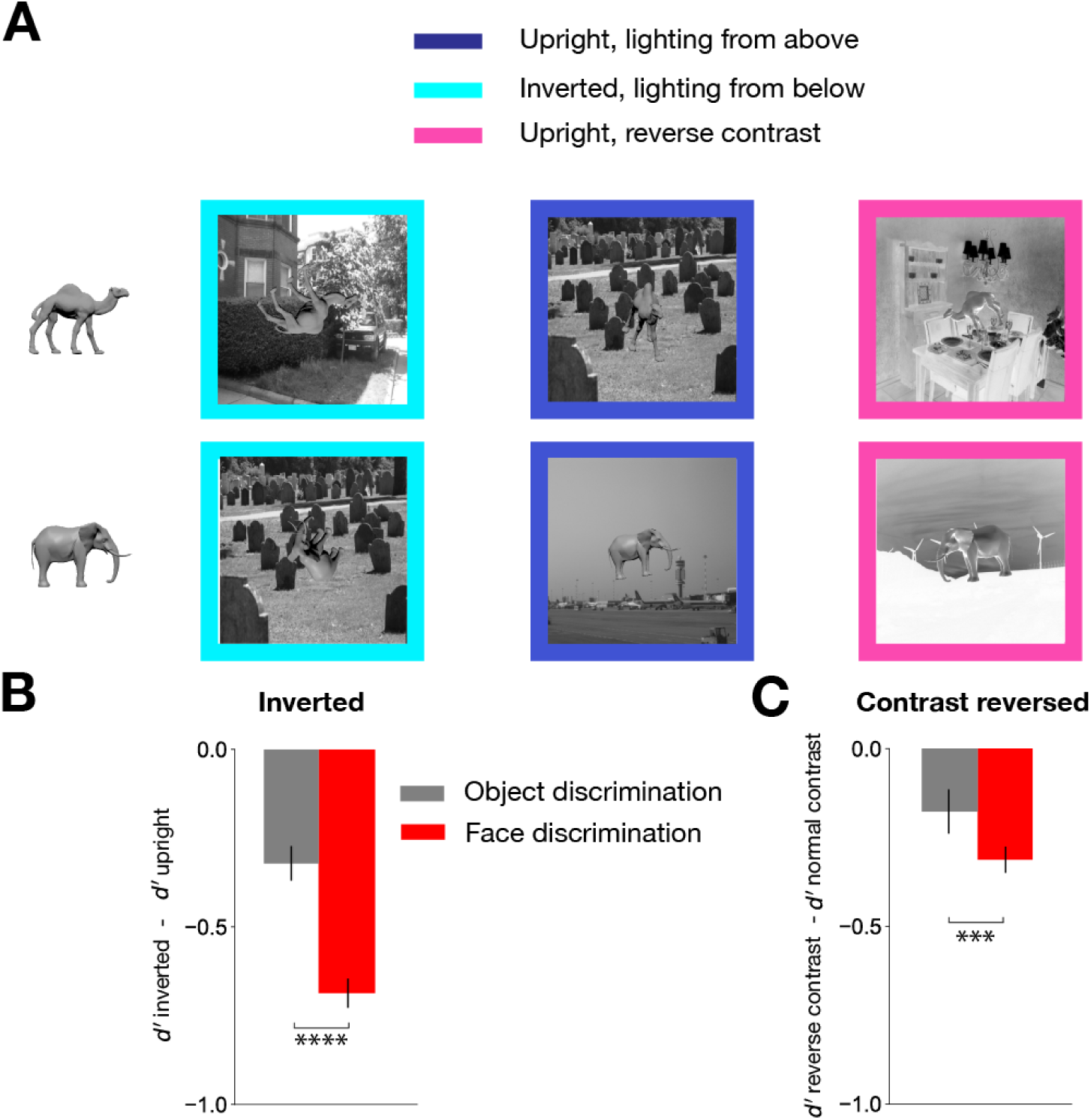
Control object discrimination task and human performance comparison to face discrimination task. **A *Object stimuli***. Token camel and elephant images. Same stimulus generation process as in the face discrimination task was used. An example image from the inverted, upright, and contrast reversed condition is shown in order. Upright/normal contrast condition only included lighting from above, inverted condition only included lighting from below, and contrast reversal condition only included upright objects in order to remove any interaction effect between conditions. Each condition included 50 images in total, 25 from each object category. **B *Inversion effect*.** Humans showed greater inversion effect for the faces than for the nonface objects. We randomly sampled 25 images from each context, upright and inverted, out of 50 images and calculated the difference of the average *d’* between the two. We repeated this process several times to obtain the error bars, which indicate 95% CI. **C *Reverse contrast effect*.** Humans showed a greater reverse contrast effect for the faces than for the objects. Error bars were calculated the same way as in **B**. Here, we randomly sampled 25 images from reverse contrast and upright condition, respectively.

### Face inversion and contrast reversal deficits in marmosets

Like humans, marmosets were worse at recognizing faces that were inverted, contrast reversed, or lit from below compared to standard faces which were upright, normal contrast polarity, and naturally lit from above. The inversion effect was quantitatively stronger in humans than in marmosets, while the reverse contrast effect was quantitatively stronger in marmosets (**Figure 4A**). It is important to note that our paradigm was able to expose inversion effects in marmosets even though they were trained on both upright and inverted faces (see **Methods**), suggesting the overall robustness of face inversion deficits in this task. To check whether the inversion effect in marmosets was weaker for non-face objects than for faces, we re-analyzed our previously reported marmoset core object recognition dataset and found that on the animal discrimination task used in our prior work (camel vs. rhinoceros), marmosets only showed a very mild inversion effect (Δd’_face_ = −0.31 vs. Δd’_object_ < −0.1; **Figure S3**)^10^.

**Figure 4.**
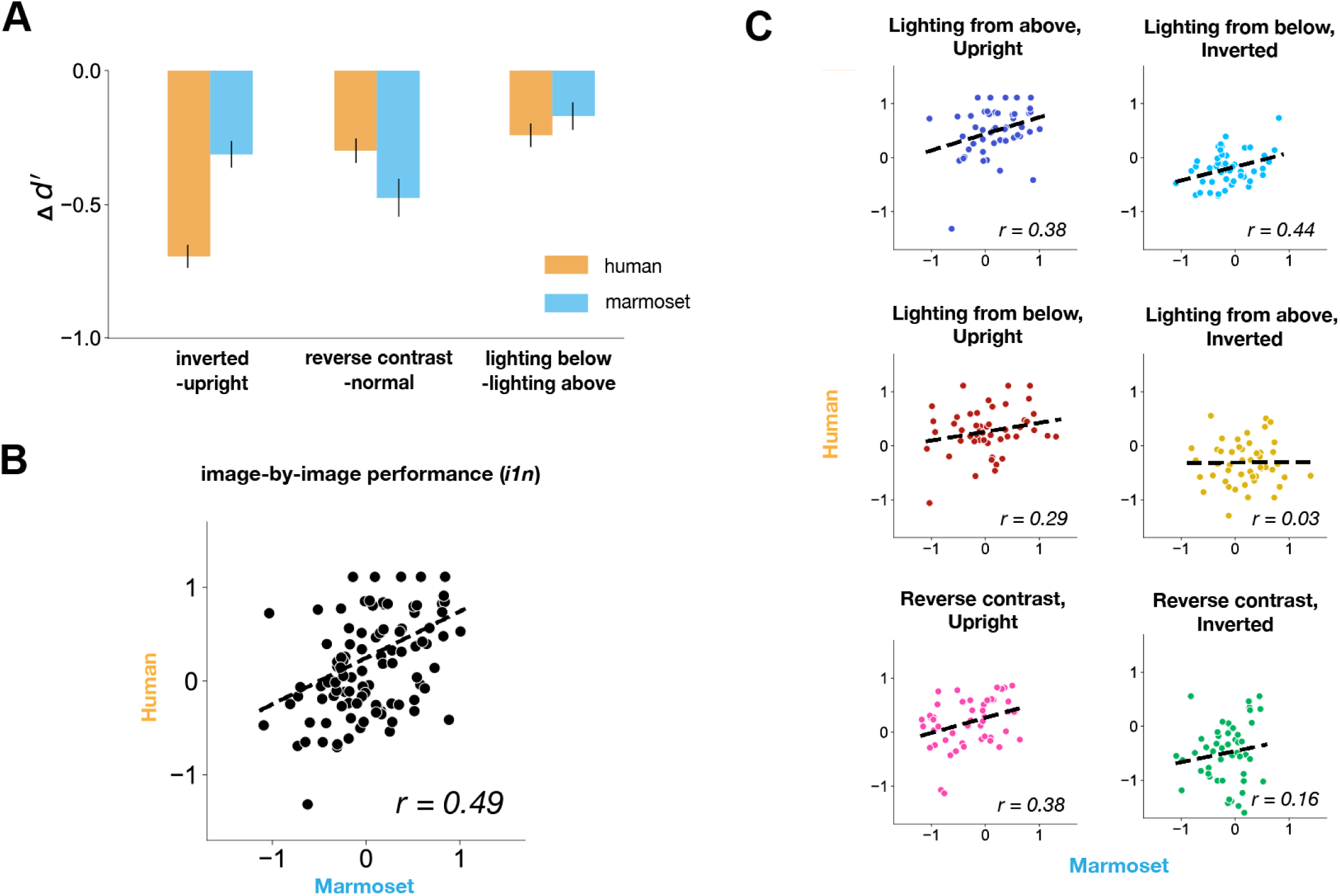
Human and marmoset image-by-image performance. A *Comparison of the effect of different conditions in human and marmoset.* Both human and marmoset subject pools showed deficits in recognizing faces that are inverted, contrast reversed, or lit from below. As in Figure 3B, upright condition only included lighting from above, inverted condition only included lighting from below, and contrast reversal condition only included upright faces. Each condition contained 50 images, 25 per identity. Similarly to **Figures 3B & 3C**, the error bars indicate 95% CI, which were calculated by bootstrapping images in each condition. B *i1n correlation.* Only considering the two congruent lighting conditions (lighting from above, upright and lighting from below, inverted), human and marmoset image-level patterns (*i1n*) were positively correlated (r = 0.49) C *i1n correlation per condition. i1n* correlations were heterogeneous across conditions, with the contexts deviating from standard, natural viewing (lit from above and inverted; contrast reversed and inverted) having the smallest correlation (right panel in middle and bottom rows). All correlation-coefficients were noise-corrected. The scatterplot for the congruent lighting condition in **B** was created by combining data from the contexts in the top row.

### Marmoset versus human behavior at the image level

To test whether marmosets matched humans at finer behavioral scales, we compared their image-by-image performance (vector of d’ per image or *i1n*, see **Methods**), which can expose differences and similarities across images that average task performance cannot^9^. Examining the natural upright and inverted conditions with congruent lighting direction, which are typically used in studies, we found a significant correlation (r = 0.49, **Figure 4B**, right panel). Further breaking out into the six individual contexts, the natural upright, lit from above condition and the true inverted condition (inverted and lit from below) indeed each separately had the highest correlations (r = 0.38 & r=0.44, respectively; first row of **Figure 4C**). However, conditions where the lighting context and the in-plane rotation created unnatural scenarios had the weakest marmoset-to-human correlation (r = 0.03 for inverted, lit from above and r = 0.15 for inverted, reverse contrast). These results suggest that marmosets and humans have more similar image-level patterns for the “natural” condition, where the light source is overhead, compared to reverse contrast faces or lighting from below the head.

## DISCUSSION

We tested humans’ and marmosets’ ability to recognize textureless faces in a fine geometry discrimination task that was not easily performed by standard feedforward DNNs. We found that both marmosets and humans could robustly recognize face identity when only given shape cues, and that their face recognition performance was at a similar level to their object recognition performance, unlike for artificial vision system controls. Moreover, humans and marmosets shared face-specific perceptual deficits due to inversion, contrast reversal, and changes in the lighting direction. Thus, high-level face recognition capabilities are shared across disparate simian primate species (New World to Old World, small to large), potentially reflecting conserved high-level visual cortical areas.

### Species comparisons across visual tasks

Our study indicates that humans and marmosets share coarse-level similarities in recognizing faces, but diverge at the image-level especially in certain lighting contexts. This result is in contrast with our previous finding that marmosets are similar to both humans and macaques in their fine-grained patterns of core object recognition (noise-corrected r ∼= 0.73)^10^. We first note that the prior object recognition task did not include changes in lighting direction and reverse contrast, which to some degree introduced image-level divergences across marmosets and humans on the face task (**Figure 3C**). Without these different conditions, image-level correlation was significantly higher (r=0.49) (**Figure 3B**). Nonetheless, further experiments could probe the marmoset-human comparison on face recognition in greater detail: (1) running each context as a separate task more like prior work in object recognition to ensure a consistent task strategy and (2) compare face recognition signatures on conspecific (marmoset faces) versus the human face recognition task used here. Previous studies have reported mixed results with regards to the species-specificity of inversion effects^35,36^. Chimpanzees exhibited the inversion effect only for chimpanzee faces, while rhesus macaques showed the inversion effect for both rhesus and human faces^17^. Capuchins also showed the inversion effect for both conspecific and human faces^37^, while squirrel monkeys showed an inversion effect only for human faces but not monkey faces (both conspecific and heterospecific)^16^. To compare with these studies, it will be necessary to test marmosets on marmoset faces in addition to human faces. Whether or not image-level patterns match better for the conspecific faces across species needs to be investigated. Thus far, we have shown that marmosets at least display an inversion effect for human faces, echoing the work from macaques, squirrel monkeys and capuchins, and possibly diverging from chimpanzees. We note that different task paradigms are used across studies, which might explain some of the differences.

### Marmosets as a model organism to study geometric face perception

Lingering doubts about the marmoset’s face recognition ability stemmed from their natural arboreal habitat — the tree canopy in the Amazonian rainforest — where visual access may be hindered by the thick foliage^38^. Moreover, their rich vocal repertoire is frequently suggested to be a primary means of social interaction, including identifying conspecifics in such a natural, visually occluded habitat.^39–42^ Nonetheless, in keeping with the ethological perspective, it is also important to consider that marmosets maintain tight bonds with their partners^43^, use various facial expressions^44^, and engage in cooperative breeding,^45^ activities that require close physical distance, close enough for visual face recognition. In lab settings they preferentially look at conspecific faces in photographs and use the gaze direction of other individuals to locate food^46,47^. These observations point to a possibility that marmosets are capable of identifying individuals via visual face recognition. Still, another reason for lingering skepticism is whether such a small simian primate brain could support face discrimination in challenging human-level tasks^48,49^. Building upon our previous work showing that object recognition in marmosets is a surprisingly good match to that in macaques and humans, the present study rigorously tested marmosets’ ability to recognize faces. Indeed, we intentionally chose to use human faces to test the limits of marmoset visual aptitude in performing similar tasks as humans. Our study provides critical behavioral evidence affirming that marmosets may serve as suitable model organisms to study the neuroscience of shape-based face perception. These demonstrated behavioral similarities from among the smallest monkeys to humans are consistent with the modest expansion of the ventral visual cortex across marmosets, macaques, and humans relative to the rest of the cerebral cortex and are consistent with the presence of potentially homologous face processing neural networks in these simian primate species^18,19,23,24^.

### Distinct roles of texture and shape in face processing

Human face perception research has made a distinction between the role of shape and texture cues in face processing^21^. Shape refers to the 3D geometry of the face, whereas texture arises from a combination of the pigmentation, illumination, and shading from shape (surface curvature). Texture seems to be the dominant cue for recognizing familiar faces, while both texture and shape cues are important for discriminating unfamiliar individuals^50–55^. In addition to familiar face recognition, pigmentation-based face discrimination, but not shape-based discrimination, is selectively impaired by contrast negation^56^. On the other hand, shape plays a critical role when learning a new identity or generalizing to different views^57–60^. Our study probed the marmoset’s ability to discriminate shape defined textureless faces with a particular emphasis on doing so across viewpoints and lighting.

Future studies could test the complementary role of texture cues in an invariant fine face discrimination task, including examining the strength of inversion and contrast reversal effects when adding texture cues. For example, does adding texture significantly improve performance across viewpoints, and does contrast reversal lead to bigger performance decrements when discriminating between textured faces? As tasks expand beyond identification, we anticipate an interplay between the type of task and the specific facial features that are emphasized. For example, previous studies found that chimpanzees relied on color more than shape to discriminate the age of conspecifics while Japanese monkeys used facial morphology to discriminate the sex of conspecifics^61,62^. Future work can test whether or not marmosets exhibit particular dependence on either shape or texture cues for various forms of inference in faces, and if this dependence on either cue is specific to conspecific faces.

### Fine 3D geometry discrimination exposes gap between biological and artificial systems

Our novel, geometric face discrimination task challenged a state-of-the-art ImageNet-pretrained DNN in a way that the object recognition task did not (**Figure 2B**). The prior basic-level object recognition task was made more difficult by including object position variation to thwart low-level features. Here, without requiring any position variation at all (all faces are presented centrally in the image, effectively at the fovea), we found that fine face discrimination was computationally challenging not just for low-level features but for the best layers in the network as a whole. The difficulty inherent to our fine face discrimination was created by naturally varying 3D pose and lighting rather than by using large 2D position shifts to different parts of the visual field of early model layers. Instead, overcoming local 3D variations to detect fine geometry differences proved challenging even for high-level features in artificial systems. Indeed, when we reduced the strict geometry demands of the task, the DNN tested (ResNet-50) benefitted in task performance from adding even the most minimal pigmentation – no hair, but adding skin color and texture (**Figure S4**). Future studies can investigate if making artificial systems more shape-sensitive, either by training them on similar stimuli to ours, or optimizing for reconstructing the underlying 3D meshes such as in depth estimation tasks^63,64^, can make them perform better on the fine face discrimination task and whether, as they perform better on the task, they also go on to exhibit similar image-level patterns as humans and marmosets. In this way, insights from high-level face recognition behavior, where primates have evolved to excel, can help bridge the remaining gaps between biological and artificial vision.

## METHODS

### Stimuli

#### Face stimuli

We created two identities for the face discrimination task – identity A and identity B – by utilizing Apple’s face tracking capability in iOS devices (see captured meshes in **Figure 1A**). The TrueDepth camera in Apple iPhones estimates the relative depth of points projected on a face. Developers can access the coordinates of the imaged 3D face mesh along with blendshapes – coefficients representing the detected facial expressions via movements of the estimated 1220 vertices – and other information through the Apple ARKit API (https://developer.apple.com/documentation/arkit/arkit_in_ios/content_anchors/tracking_and_visualizi <u>ng_faces</u>). For acquiring the particular face meshes used in this study, we built upon FaceCaptureX, an iPhone application written by Elisha Huang^65^, which returns the 3D mesh coordinates of the face currently in view along with the blendshapes and photos of the face at a given framerate. We edited the source code of the application to collect extra information including the projective transform and the resolution of the captured photos. After collecting a face mesh using the customized version of FaceCaptureX, the mesh was stitched onto a faceless, dummy 3D head model to generate the full 3D face plus head object using custom code written in MATLAB (code available at https://github.com/issalab/Jeon-apple_facestim_generation). The stitched 3D face mesh plus head mesh was then exported to the open-source Blender software for mesh editing (www.blender.org). Eyes were added to the full 3D face mesh, and the entire mesh was smoothed using the *Shade Smooth* function in Blender, which locally interpolates vertex normals. Critically, we added identical head and eye meshes to face meshes from both identities to minimize the difference between the two identities other than their internal face geometry.

Identity A was modeled after one of the lab members (40 yo male) and Identity B was modeled after a rhesus macaque (6 yo male), both with neutral expressions. Ultimately, we chose face meshes from different primate species to create a greater separation of face geometry. While this species geometry discrimination task might be easier, the task provided a good starting point for probing marmoset face recognition ability and proved to be surprisingly challenging for DNN model controls. Human behavior data on this task showed that the inversion effect is comparable in magnitude across the two identities (t = 0.898, p-val = 0.375; **Figure S5**), suggesting that in this identity discrimination setting, both identities engaged signature aspects of face processing.

From each identity, we rendered 151 images generated using different 3D viewing parameters (images available at https://github.com/issalab/Jeon-et-al-Marmoset-Face-Behavior and see **Figure S6**). To emulate real-life interactions with faces where humans usually foveate on an individual’s face via centering eye movements, we consistently placed the 3D foreground face centrally on top of a randomly chosen natural image background. Ten backgrounds were randomly selected from an open-source database and used across both identities such that the background identity provided no information about foreground face identity^66^. In each image, the face was randomly scaled between ∼4.5 deg to 9 deg, and rotated horizontally (−90 to +90) and vertically (−45 to +45). All three parameters – size, horizontal rotation, and vertical rotation – were drawn randomly from the uniform distribution of their respective range. While previous research that tested object recognition in marmosets used a greater range of views for objects – −180 to +180 in all three axes – we removed the extreme poses for faces here to ensure that most of the internal facial features were visible, as opposed to viewing the back of the head. Next, we added essentially three different lighting contexts where the faces were either lit from above (i.e. normal lighting), lit from below, or contrast reversed. In each lighting condition, the faces were presented either upright or inverted. In total, there were 6 different contexts (three lighting contexts by two in-plane rotations) with 25 images per context. In the reverse contrast condition, both the face and the background were contrast-reversed to prevent performance from being affected by the ease of foreground-background segmentation. For **Figure 3** and **Figure 4**, upright condition included only the “true” upright condition, where the faces are upright and lit from above. Inverted condition only included the true inverted condition, where the faces are inverted and lit from below. The reverse contrast condition only included the upright faces, in order to remove any interaction effect between the lighting context and in-plane rotation context. These three conditions are colored navy, cyan, and pink, respectively in **Figure 1**. In addition to these 150 images per identity where the face was overlaid on a background, we included a token image with no background but containing an upright, downlit face in the frontal view (0° rotation) presented at ∼9 degrees size. The entire rendered image including the background subtended ∼20 degrees in the marmoset touchscreen setup in their homecage (where the marmoset head was generally positioned near the juice reward tube at a fixed distance from the screen). For humans, who performed trials on a computer, assuming typical ergonomic arrangements, we estimate that the images (face plus background) were presented at 4-12 degrees in visual angle.

For **Figure S4,** textured face meshes were created using one of the photos taken from the FaceCaptureX application. A photo was selected and edited in Photoshop to remove non-face areas of the image so that only the texture of the face area remained, and the outside was colored in a similar color as the skin of the face. This photo was then used as a texture map for the 3D face mesh. Like the textureless meshes, the textured meshes were stitched to the head mesh in MATLAB and the full face plus head mesh was exported to Blender for mesh editing. Eyes were added, the entire mesh smoothed, and the texture map applied in Blender.

All stimuli were generated using MkTurk, a web-based behavioral platform developed in the lab (<u>mkturk.com/landing</u>; code available at https://github.com/issalab/mkturk). See Methods in Kell et al. 2023 for further details about MkTurk. In addition to presenting 2D images, MkTurk is equipped with 3D scene creation capability using the open source ThreeJS library for rendering in WebGL (www.threejs.org). Experimenters can specify scene parameters, including all the viewing parameters listed above to modify 3D object renderings upon every displayed frame using the real-time web-based 3D rendering engine. All rendered image stimuli are shown in **Figure S6**.

#### Object stimuli

Two object classes – camel and elephant – were chosen for the control object discrimination task. As in the face discrimination task, these categories reflected a species discrimination task between two animals that are relatively similar in general layout – four legged, similar body plan – but different in the exact geometry. A 3D object model for each animal class was downloaded from the SketchFab library (https://sketchfab.com/). Both objects were rendered in the same gray color. We generated 151 images of each animal class using the same generative distribution of viewing parameters and applied the same contexts as in the face stimuli (including lighting direction changes, contrast reversal and inversion). For the canonical token image, we chose a side view of an upright camel and elephant, as compared to the frontal view for faces. All rendered image stimuli are shown in **Figure S7**.

### Two alternative forced choice stimulus-response task

Humans and marmosets participated in the face discrimination task, and humans were tested on the follow-up object discrimination task. Subjects initiated a trial by touching a dot at the center of the screen. A sample image was then flashed (250 ms for marmosets and 200 ms for humans). After the sample image disappeared, two token images appeared that were representative of the two identity choices, and the subjects chose which of the two identities was seen in the sample image. The choice screen was always fixed such that the token image of Identity A appeared on the left, and Identity B on the right, making this a stimulus-response task (i.e. always choose left for identity A). For the object discrimination task, camel always appeared on the left and elephant on the right.

### Marmoset behavior

Four common marmosets (*Callithrix jacchus*; subjects J, G, V, and S for AJ, Bourgeois, Bolshevik, and Sausage) were tested on the same face discrimination task. All procedures were in compliance with the NIH guidelines and the Columbia University Institutional Animal Care and Use Committee (IACUC). Marmoset behavioral data was collected in their homecage, where an individual monkey performed trials on a touchscreen device (Google Pixel C tablet or Google Pixel 4xl phone). Marmosets in their homecage participated in the task about 3 hours per day for a juice reward. For testing on the full task, monkey J, G, and V completed ∼1800 trials per day for ∼3 months. Monkey S completed around 600 trials per day. In total, Monkey J finished 106,717 trials, Monkey V 189,988 trials, Monkey G 149,438 trials, and Monkey S 2,843 trials. Subjects were given up to 5 seconds to make a choice, and all data presented in this paper, both individual and pooled, used trials where the monkeys made a choice (left or right) between the two identities within the five second choice timeout period. All four monkeys went through a series of training stages. For the first stage, they discriminated between the token images with no pose variation and no backgrounds. We slowly added more variations such as size, pose, position, and background. In the final stage of training, the marmosets were introduced to the high-variation images that were similar in complexity to the actual test images. Once the monkeys saturated their performance on training images, we switched them to the testing image set for the face discrimination task. Lapse rate for the marmosets was calculated by partitioning the trials into two halves, finding the best-performing image in one half, and calculating the cross-validated performance for that image in the other half of trials. For all main analyses of the paper, we pooled trials across the four marmoset subjects.

### Human behavior

Human behavioral data were collected from the Amazon Mechanical Turk platform (MTurk), where subjects participated either in the face or object discrimination paradigm described above. All data were collected in accordance with the Institutional Review Board of Columbia University Medical Center. For the face discrimination task, 37 subjects were recruited to complete ∼500 trials per subject. Seven of them were removed from the subsequent data analysis because they either performed near chance (0.5) or completed fewer than 20 trials. While Amazon MTurk provided a convenient way to amass a large number of psychophysics trials, it can be uncontrolled relative to a lab setting and thus, might include subjects that are less motivated and focused. A large performance spread on the face recognition task in the 30 subjects (76.57 ± 12.09%) indicated potential non-perceptual variability in task performance. Thus, we recruited 4 out of the 30 subjects to collect a larger number of trials per subject (8,011.5 ± 418.9 trials), more on par with the number of trials per marmoset subject. Indeed, the 4 subjects performed much better than the 30 subjects, at around 91% (**Figure 2B**). A much larger number of trials from a smaller number of subjects also enabled calculating a more reliable image-by-image difficulty in the face task (i.e., higher internal consistency of data on *i1n*, see **Methods** “quantification metrics and statistical analysis”). We pooled trials across the 4 subjects for all analyses. Lapse rate for the face discrimination task was calculated in the same way as for marmosets. For **Figure 1D**, we bootstrap resampled trials from the 4 subjects 1000 times and reported the mean and the standard deviation of the performance. For the object recognition task, 31 subjects were recruited to complete ∼500 trials per subject. Four of them were removed from the data analysis following the same exclusion criteria as in the face discrimination. As in the face discrimination task, we pooled trials across the 27 subjects for all analyses.

### DNN model behavior

We used the off-the-shelf Resnet-50^25^ and VGG-face^29^ pretrained, convolutional feedforward models for DNN model controls. Resnet-50 is a state-of-the-art deep net trained on the ImageNet dataset and excels in object recognition from natural images. By contrast, VGG-face is specifically trained on a large-scale human face dataset to individuate identities across a large corpus of face photographs. For each model, we trained linear classifiers on the activations from the penultimate layers and reported the classifier performance (see Kell et al., 2023 for details). To train linear decoders, we created a training dataset that was distinct from the actual test set but used similar distributions for latent variables – rotation, size, lighting, and background – as the test set for measuring human and marmoset behavior. We used the same uniform distribution between ∼4.5 deg to 9 deg for size, −90 to +90 for horizontal rotation, and −45 to +45 for vertical rotation. For lighting, we randomly varied the lighting direction horizontally from −1 to 1 (left to right), and vertically from −1 to 1 (below to above) in order to expose the models to more lighting variations than there exist in the test set, which only included two lighting directions (directly above or below). We did not include reverse contrast images in the training since this condition is not part of our regular experiences with faces. We ran the images of the training and test sets through the network, and obtained the activations from the penultimate layer of that network (2048 activation units per image). We trained linear classifiers with these activations for different numbers of images in this training set. We randomly sampled different training images 1000 times for training linear support vector machines (SVM) with L_2_ regularization and a hinge loss. For the main figures in the paper, we chose the number of images that saturated performance on the test set (approximately 800 images per class, see **Figure S1**). Model performance was reported as the mean and standard deviation of the classifier performance over the 1000 iterations of resampling images for classifier training. For each image, d’ was calculated as the distance from the hyperplane learned by a linear SVM.

### Behavioral performance metrics and statistical analysis

For each individual image *j*, behavioral *d’* of primates was calculated as follows:

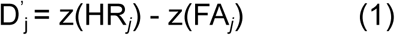

Where *HR_j_* and *FA_j_* are the hit rate and the false alarm rate – proportion of trials where the image *j* was correctly classified and the proportion of trials where any image was incorrectly classified as the object class that the image *j* belonged to, respectively. *i1* is a vector of d’ for all the images for a given discrimination task. Thus in this study, *i1* is a 302-length vector that contains a *d’* estimate for the 151 images from each of the 2 classes. The normalized *i1*, or *i1n*, was calculated by subtracting the mean performance for the correct class (average d’ across 151 images) from each d’*_j_* to remove any mean difference in difficulty between the two classes, and the figures and texts refer to this class mean subtracted *d’* as *i1n*. After calculating *i1n* of the pooled human and pooled marmosets, these image-by-image performance patterns were compared against each other by calculating the noise-corrected Pearson’s correlation coefficient. The coefficient was calculated by randomly partitioning the trials into two halves, computing the *i1n* for each half, and calculating the mean of the correlation of the human and marmoset *i1n* across the two split halves. We normalized this split-half correlation by the internal consistency of *i1n* within each species. The internal consistency was calculated as the geometric mean of the correlation coefficients of the *i1ns* across the two trial split halves within each species (to provide a ceiling correlation given trial-by-trial noise in the data).

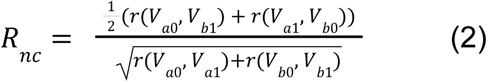

Where r() calculates the Pearson’s correlation coefficient, and *V*_*a*0_, *V*_*a*1_ refers to two split-halves of a system *a*, and *V*_*b*0_, *V*_*b*1_ to two split-halves of a system *b*. This split-halves noise ceiling was used to normalize raw correlation values to produce the noise-corrected correlations reported in **Figure 4B and C**.

## ACKNOWLEDGEMENTS

The authors thank Elizabeth Yoo and Sophie Bokor for early project support, and Anthony Lee Williams and Sarah Latoya Wilson for technical support. This work was performed on the Columbia Zuckerman Institute Axon GPU cluster. EBI was supported by a Klingenstein-Simons fellowship, Sloan Foundation fellowship, and Grossman-Kavli Scholar Award. The authors declare no competing interests.

---

**Figure S1.**
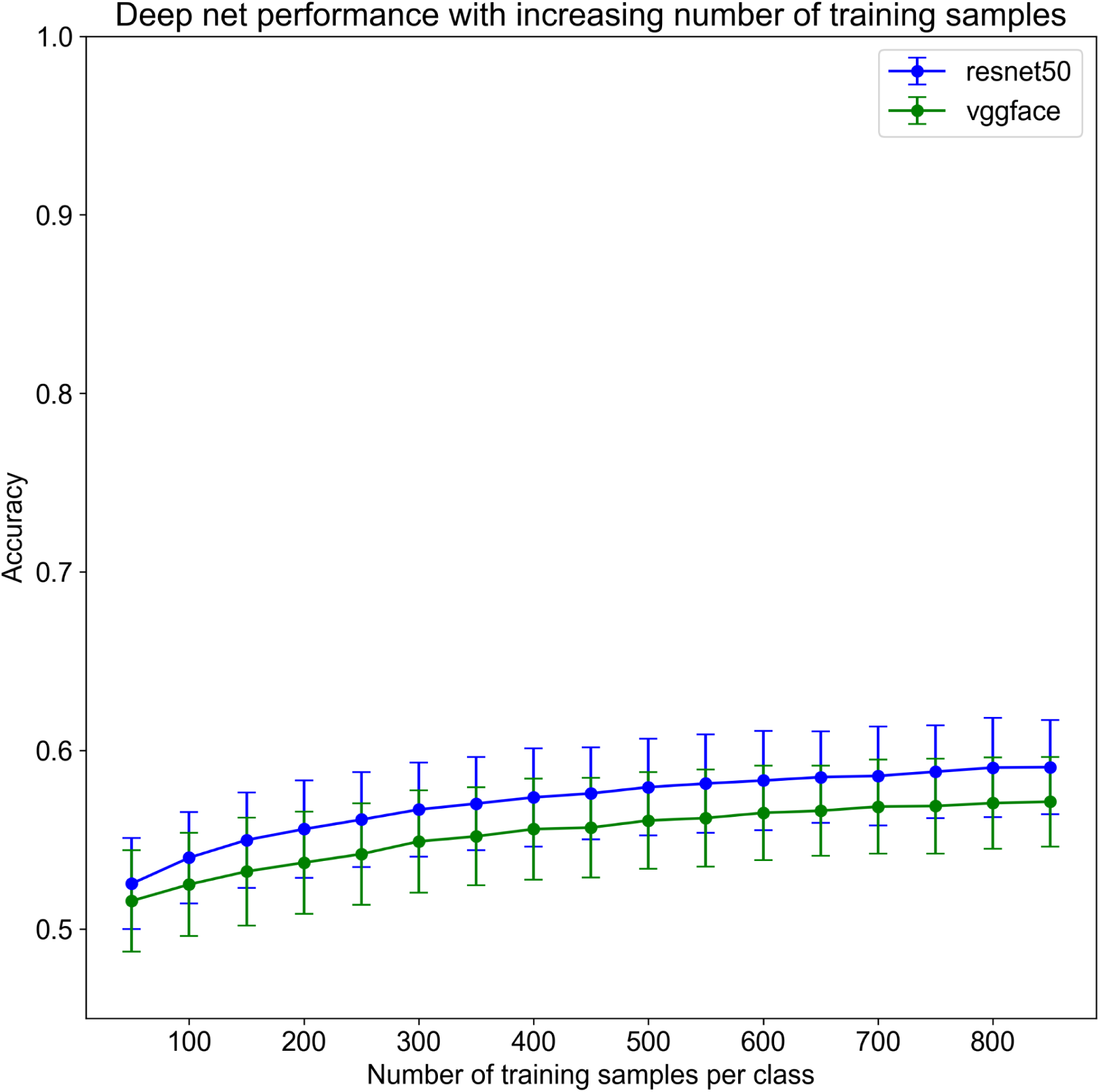
DNN performance on the face discrimination task as a function of number of training images. In order to ensure that we reached a performance plateau with the deep net controls, we trained linear decoders with different numbers of images per identity in 100 image increments. Bootstrapped error bars are standard deviations over 1000 classifiers trained on random samples of training images. The performance number from using 800 images/identity in classifier training was used in Figure 2B in the Main Text.

**Figure S2.**
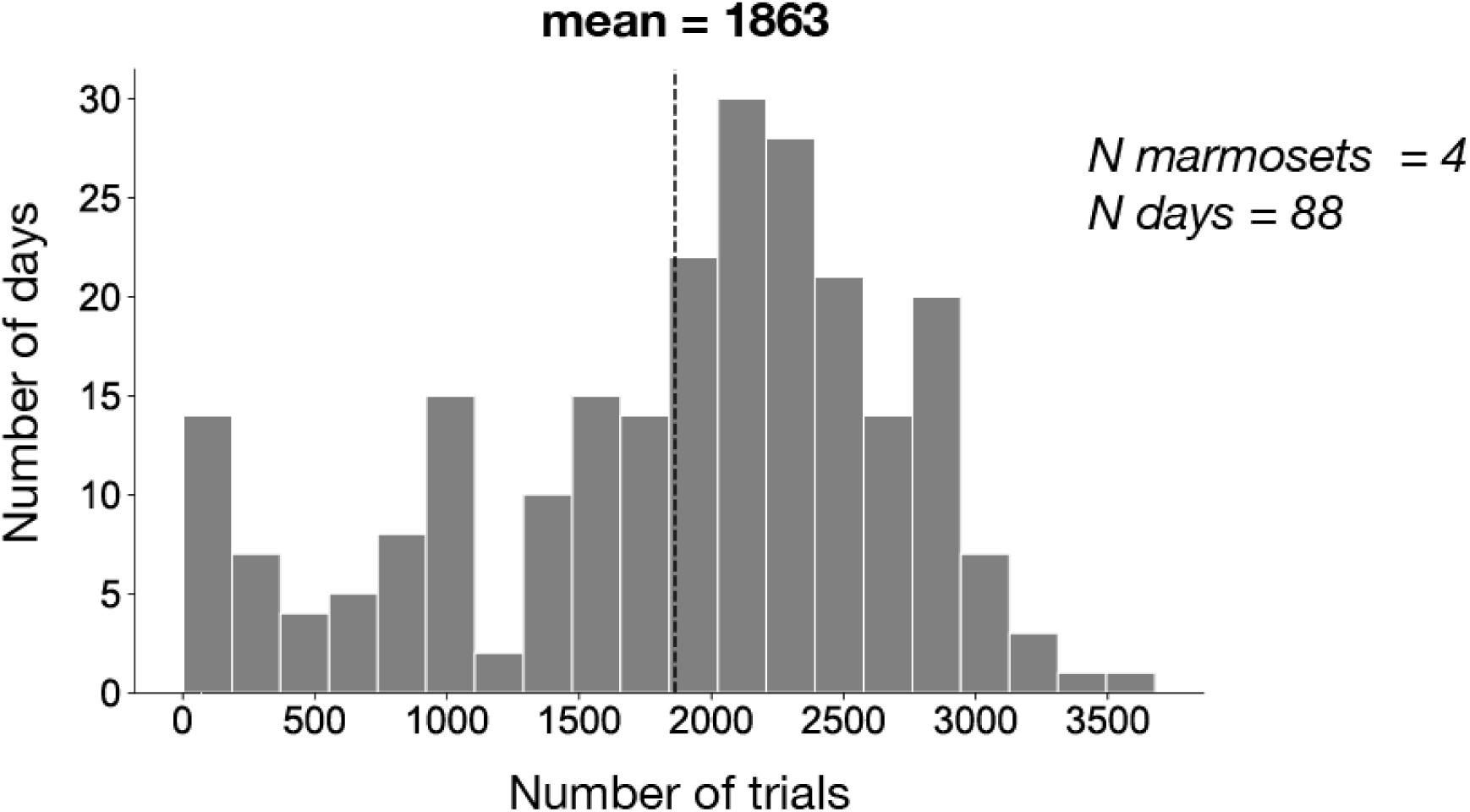
Number of trials performed per day by marmosets. Three monkeys were tested over a period of 2-3 months. The fourth monkey was tested over a period of a week.

**Figure S3.**
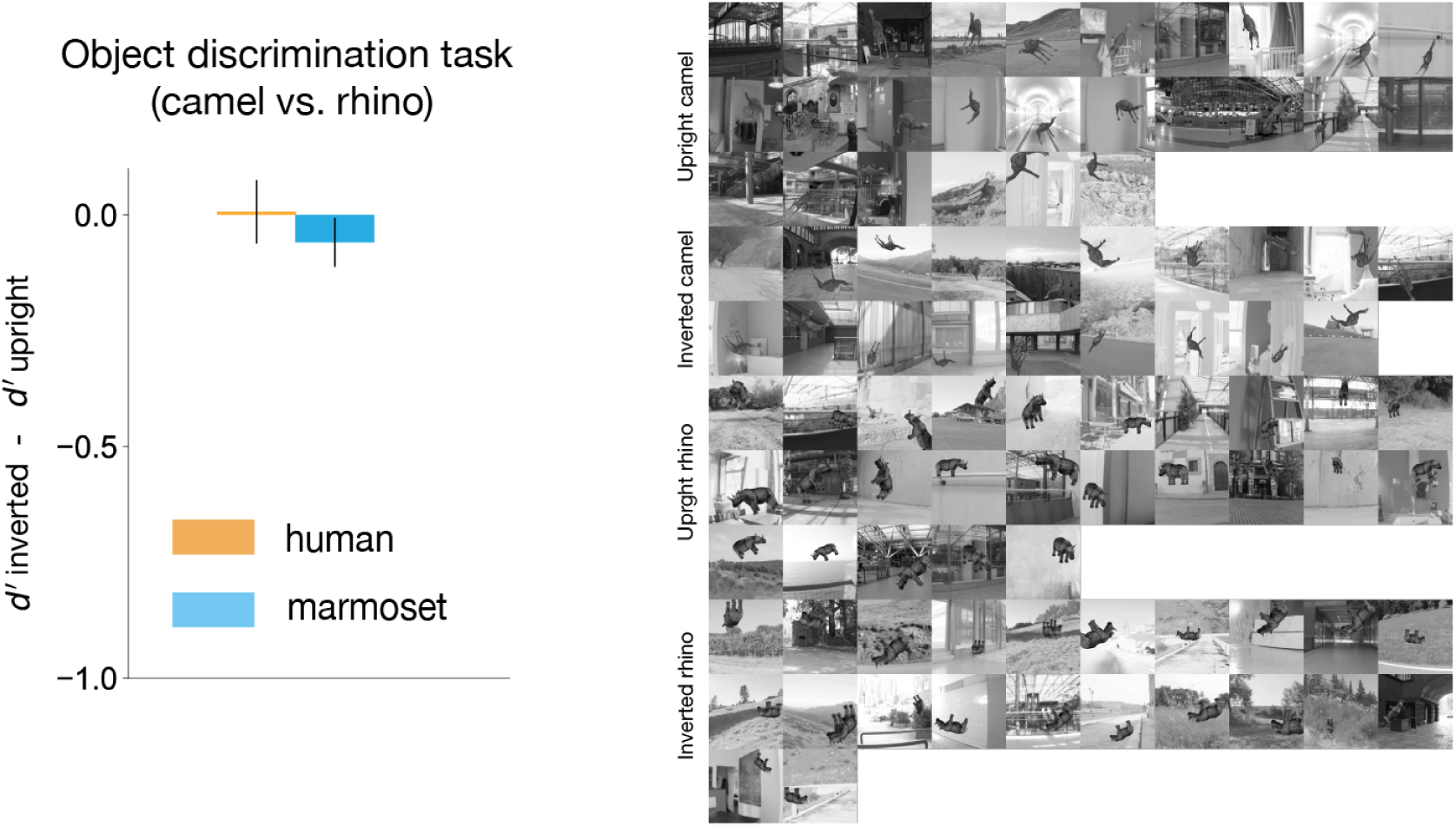
Inversion effect in an object discrimination task (camel vs. rhino). Re-analyzing data from Kell et al. 2023, we found that marmosets suffered a very mild inversion effect (Δd’ < −0.1) in a camel vs. rhino discrimination task. In this task, 100 images from each object category were presented to subjects. Among these, 26 images of upright camel, 19 images of inverted camel, 25 images of upright rhino, and 22 images of inverted rhino were chosen. We randomly sampled 20 images from each condition, upright and inverted, and calculated the difference of the average *d’* between the two. We repeated this process 50 times. Error bars indicate 95% CI over the 50 repeats. All stimuli used in the analysis are shown on the right.

**Figure S4.**
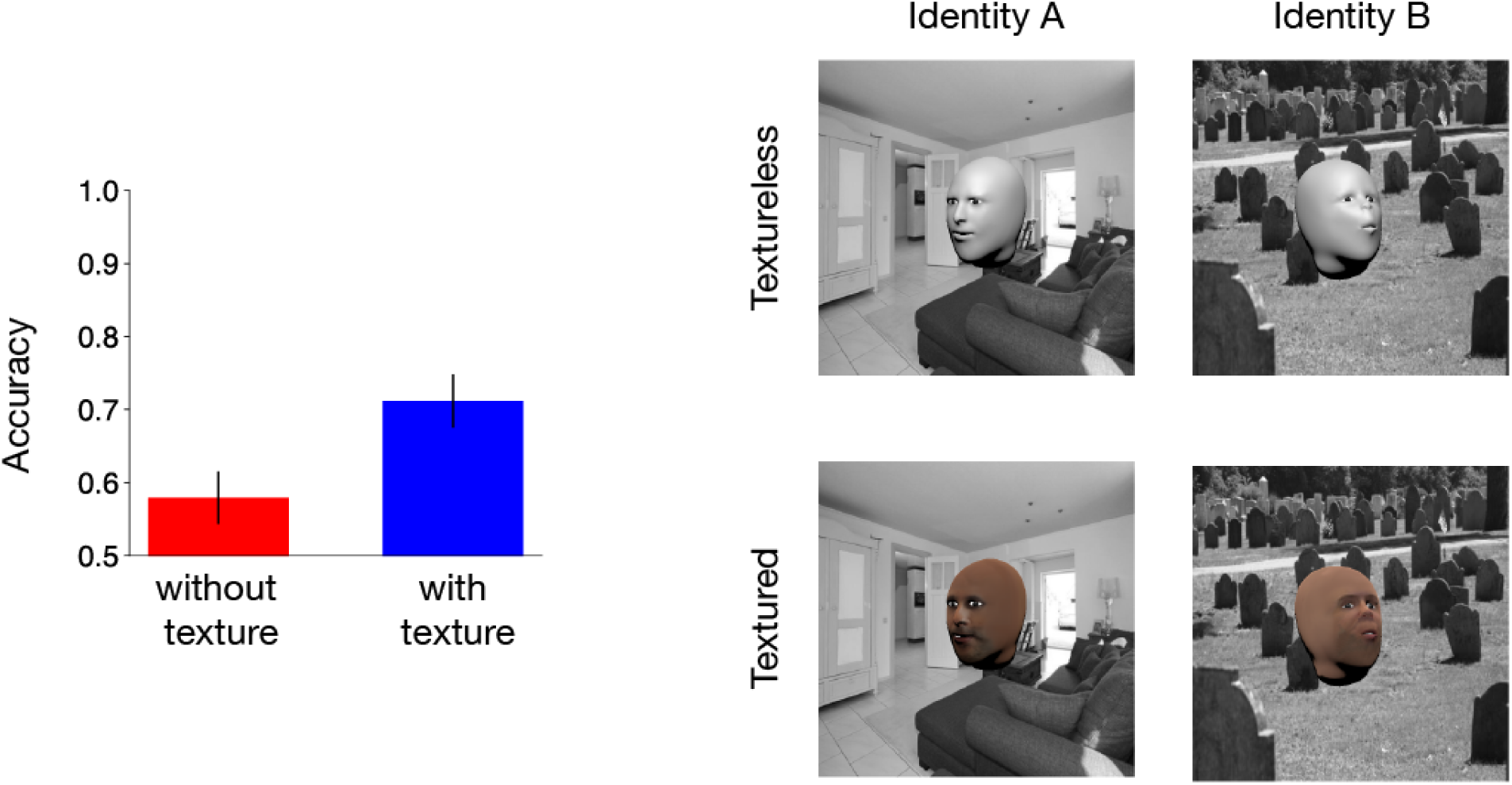
Comparison of ResNet-50 performance for textured vs. textureless face discrimination. When texture is added to the face, model accuracy on face discrimination significantly improves. An example image from textureless and textured versions of each face identity is shown (right panel).

**Figure S5.**
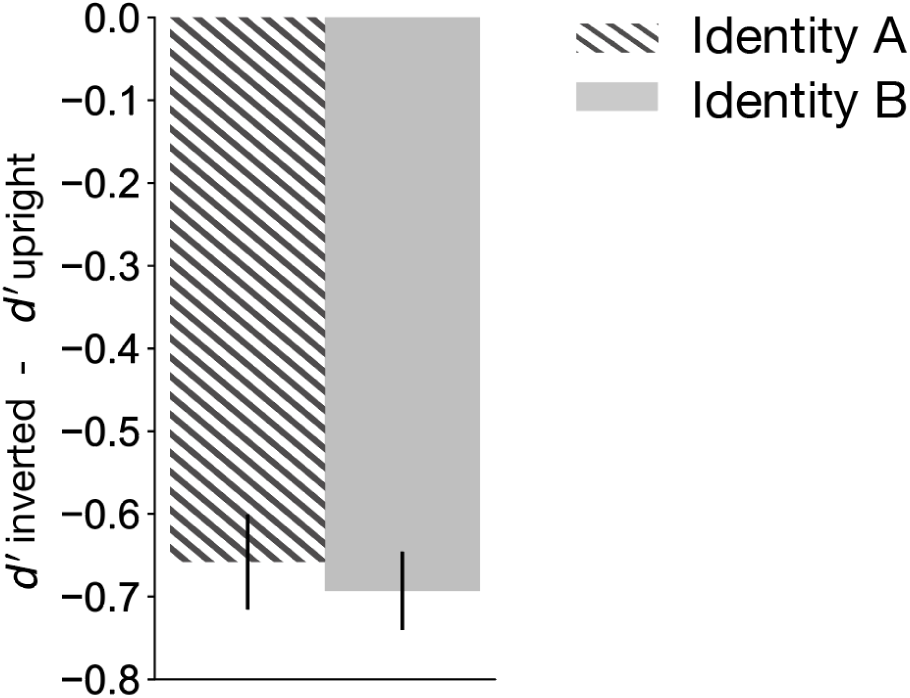
Comparison of inversion effect for two identities. In order to establish that both identities evoked face-specific perceptual effects, and at a comparable level, we quantified the inversion effect in human subjects per identity and found that both Identity A and Identity B induced a similar level of inversion effect.

**Figure S6.**
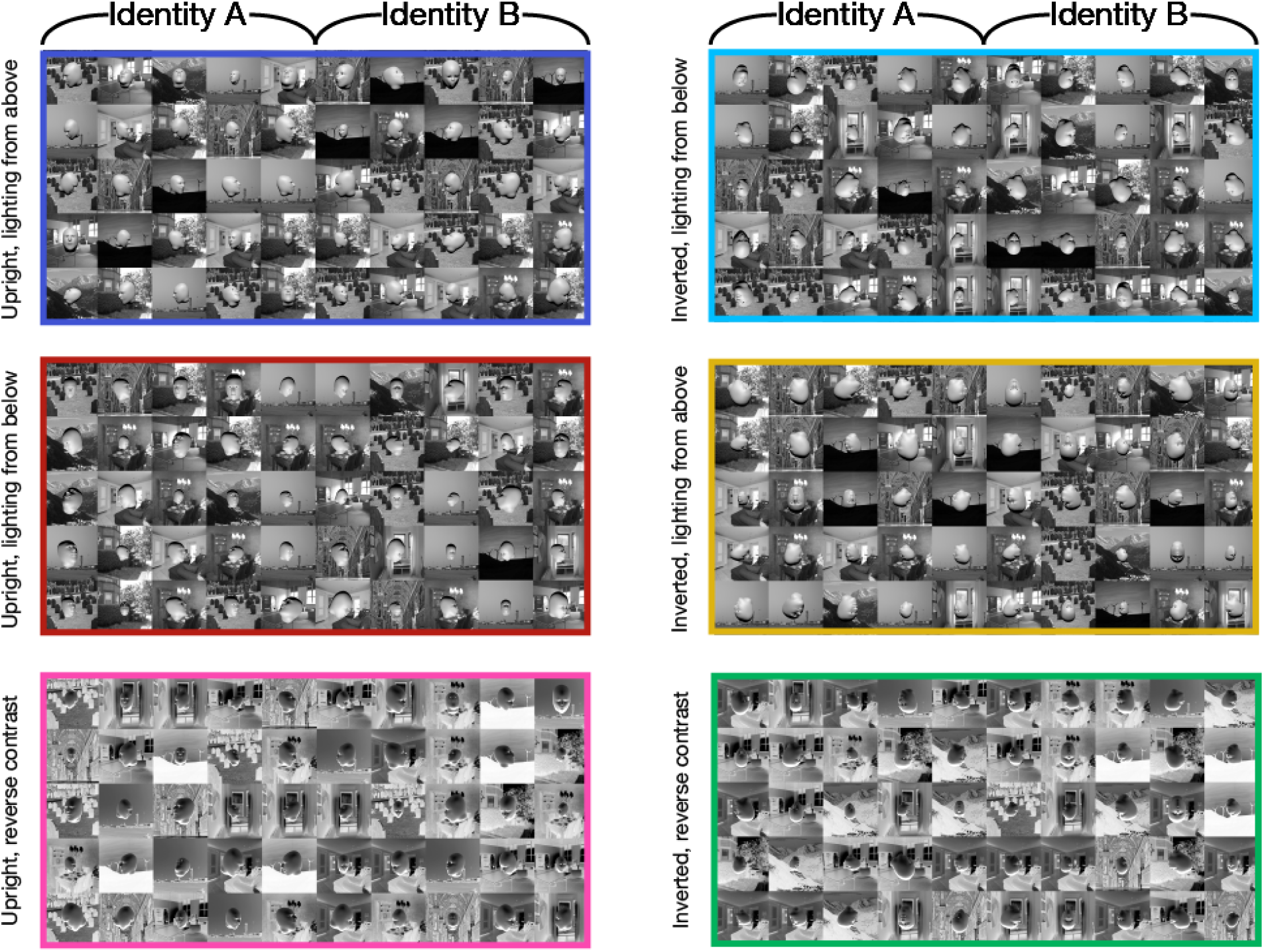
All 300 images used in the shape based face discrimination task. Different colors indicate different viewing contexts, as in Main Text Figure 1B.

**Figure S7.**
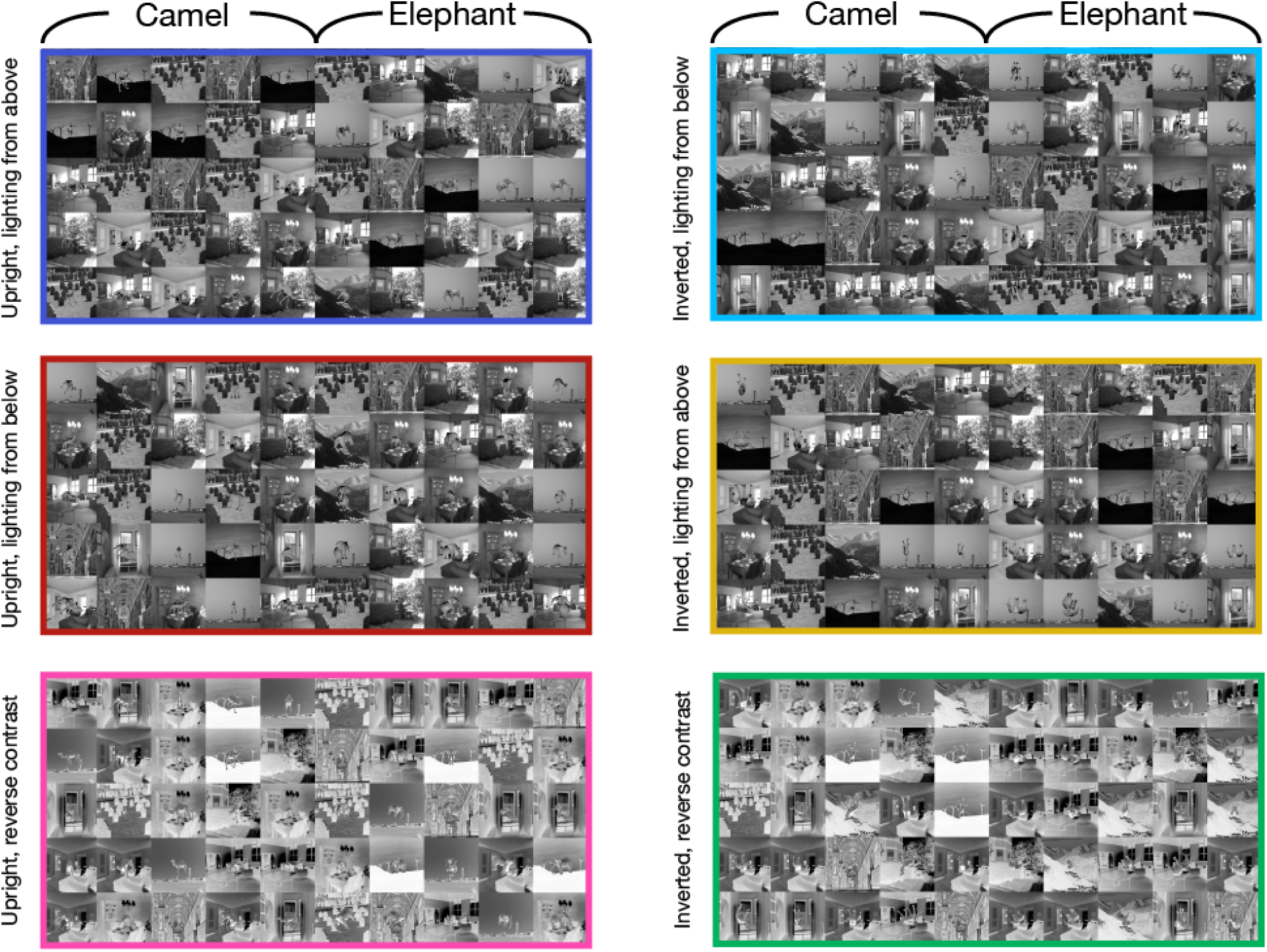
All 300 images used in the shape based object discrimination task (camel vs. elephant). Different colors indicate different viewing contexts, as in Main Text Figure 1B.

**Video S1.**
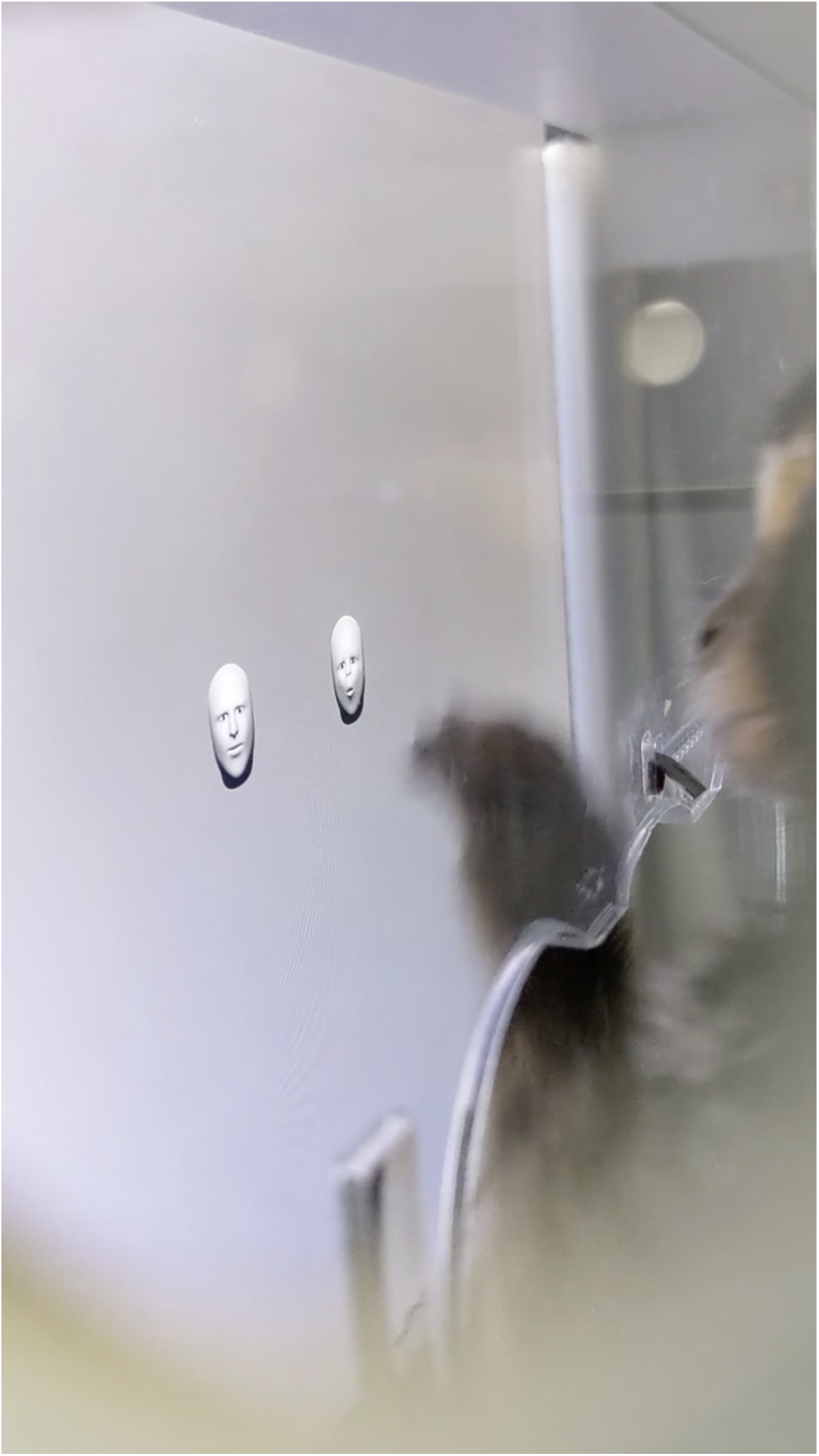
A marmoset subject performing an early version of the face discrimination task on a touchscreen device in their homecage.

## Notes

### Competing Interest Statement

The authors have declared no competing interest.

